# Directing the Chondro-Fibro Axis via Early Microenvironmental Interactions to Enable Precise and Volumetric Cartilage Repair

**DOI:** 10.64898/2026.08.13.744318

**Authors:** Maddie Hasson, Hanna Solomon, Samir Chihab, Aniya Hartzler, Lorenzo M. Fernandes, Alan Zhao, William X. Patton, Nyssa M. Morgan, Alan Y. Liu, Nazir M. Khan, Jarred M. Kaiser, Jason T. Bariteau, Jay M. Patel

**Affiliations:** Department of Orthopaedics, Emory University School of Medicine, Atlanta GA; Joseph Maxwell Cleland Atlanta VA Medical Center, Decatur GA; Optics 11 Life Inc., Boston MA

**Author notes:** Address Correspondence to: Jay Milan Patel, Assistant Professor, Department of Orthopaedics, Emory University School of Medicine.

**Keywords:** Cartilage repair, chondrogenesis, fibrosis, cartilage regeneration, fibrin, microenvironment, microfracture, marrow stimulation

## Abstract

Successful cartilage repair remains one of the most significant challenges in the musculoskeletal field. Microfracture (MFx), a form of marrow stimulation, remains the predominant repair technique, but it exhibits routine failure due to inadequate defect fill and inferior fibrotic tissue formation. Whereas current strategies focus on augmenting MFx with scaffolds and bioactive factors, the potential to target the MFx clot itself and use the capabilities of this dynamic environment to guide MFx repair remains largely unexplored. We verified that MFx contraction and fibrosis hinder repair success in minipigs and become evident as early as one week in multiple animal models. Therefore, our objective was to investigate and direct microenvironmental interactions in the MFx clot to promote volumetric maintenance and reprogram cells from a fibrotic to more chondrogenic phenotype. Extracellular control of cell-environment interactions, through fibrinogen augmentation or anti-fibrinolytic treatment, limited contraction but had no effect on or even exacerbated the fibrotic susceptibility of marrow-derived cells (MDCs). Intracellular control of microenvironmental interactions, through modulation of the Rho-ROCK pathway, drove TGF-β3 activity of MDCs along a “chondro-fibro axis”. In particular, treatment with the ROCK inhibitor Fasudil drove TGF-β3-treated cells away from a myofibroblast phenotype and towards chondrogenesis. Short-term Fasudil treatment prevented TGF-β3-driven macroscale clot contraction and enhanced cartilage-specific matrix deposition *in vitro*. In a pilot rat study, this combination treatment improved GAG deposition and better protected surrounding cartilage. These findings suggest that Rho-ROCK modulates TGF-β signaling along this chondro-fibro axis and its precise control could be the key to promoting precise and volumetric cartilage repair through microenvironmental interactions.

## Introduction

Articular cartilage is a unique tissue with a dense composition of type-II collagen and proteoglycans that are critical to its resilience. Due to the complexities of joint motion, cartilage is often injured, initiating a deteriorative cascade. Given the low cellularity and avascular nature of cartilage, it has a limited self-regenerative capacity^1,2^, motivating the need for reparative techniques. Microfracture (MFx) is the predominant reparative approach for cartilage injuries^3,4^, and it involves puncturing the subchondral bone to recruit regenerative bone marrow elements, including autologous mesenchymal stromal cells ^5,6^. This provisional fibrin-rich marrow clot is typically remodeled into a mechanically inferior, scar-like fibrotic tissue rather than durable hyaline cartilage. Furthermore, MFx frequently results in suboptimal defect fill, which correlates with poor patient outcomes^7,8^. The increased susceptibility to wear of the MFx scar and excess stress placed on the surrounding cartilage result in >30% failure by 5 years^9^, a rate that is considerably higher in physically active populations (military, athlete). These failures necessitate reoperations and accelerate arthritis progression, motivating regenerative approaches to provide functional tissue restoration. While other therapies (e.g., chondrocyte implantation) demonstrate slight improvement over MFx, the relative costs (>$60k) and need for a second procedure (harvest, implantation) are prohibitive^10,11^. Due to procedural ease and low cost, MFx remains the gold standard of cartilage repair^12^, and similar marrow stimulation is now employed in the repair of several other musculoskeletal tissues (e.g., rotator cuff^13,14^, enthesis^15^). While a variety of scaffolds^16,17^, growth factors^18,19^, and cell types have been investigated for decades to improve MFx repair, functional and successful cartilage repair remains elusive. Thus, informed modalities to redirect these marrow-mediated repairs from inadequate fibrous scars towards volumetric functional cartilage are needed.

Both clinical and preclinical studies have advanced approaches to improve MFx outcomes. For example, autologous matrix-induced chondrogenesis (AMIC) involves placing a type-I/III collagen membrane over the MFx, leading to improved patient outcomes^20,21^ but still inadequate defect fill^22^. Other materials have included biologic (collagen, hyaluronic acid ^17,23^) and synthetic (e.g., polycaprolactone ^24^) polymers, though fixation and retention of both AMIC and these scaffolds remains a challenge^25^. To further improve healing, approaches have also incorporated chondrogenic factors, most notably transforming growth factor beta 3 (TGF-β3 ^19,23^). Nevertheless, TGF-β3 exhibits a context- and cell-dependent effect ^26,27^, with data implicating it in both tissue contraction and fibrosis, potentially exacerbating MFx failures. To our knowledge, MFx augmentation approaches have not fully considered the MFx for what it is – a rapidly contracting and fibrosis-susceptible clot – even though these early tendencies create obstacles to volumetric fill, chondrogenesis, and functional cartilage repair.

Therefore, the objective of this study was to investigate contraction (“loss of defect fill”) and fibrosis in the early MFx environment to determine how microenvironmental perturbations may prevent these problematic features. First, we established variability in MFx outcomes in a minipig model, highlighting a zone of “ideal cartilage repair” with >80% defect fill, histological tissue quality, and protection of the surrounding cartilage. In multiple animal models, we observed that this loss of defect fill and fibrotic susceptibility manifest as early as one week, motivating augmentations to address this phase. We then simulated MFx *in vitro* with a fibrin gel model with encapsulated marrow-derived cells (MDCs). We first battled contraction extracellularly through fibrinogen supplementation or anti-fibrinolytic treatment; however, these perturbations seemed to drive cell behavior towards fibrosis instead of chondrogenesis. Next, we focused on intracellular contraction control via the Rho-ROCK pathway. We found that ROCK inhibition with Fasudil mitigated TGF-β3 mediated contraction while directing marrow cells along a “chondro-fibro axis” towards chondrogenesis. We then evaluated this combination (Fasudil + TGF-β3) as a pharmacological, point-of-care augmentation to MFx, both *in vitro* and *in vivo*, demonstrating improved cartilage repair capacity. Altogether, by driving early volumetric maintenance and the “chondro-fibro axis”, our study presents a promising translational strategy for improving marrow-mediated cartilage repair.

## Results

### MFx volumetric loss and fibrosis susceptibility are critical barriers that manifest as early as one week

MFx provides short-term symptom relief and return-to-activity; yet inadequate repair tissue volume and improper tissue formation (fibrosis) prevent long-term success. Clinical studies note poor fill in ∼57% of patients^7^, and only 25% of these exhibit symptomatic improvement^28^. The reported clinical cutoff for “successful” defect fill is between 50 and 75%^29,30^. In our prior MFx study in a 12-week Yucatan minipig model^17^, we observed variable defect fill, ranging from 60-95% (**Figure 1A/B**). We also observed variability in histological quality, often with inferior fibrous tissue. From this study that evaluated scaffold-augmented MFx, defect fill was plotted versus ICRS II histological score^31^ (**Figure 1C, Figure S1**), and a significant correlation (*R*^2^*=0.37, p=0.003*) was observed, suggesting that maintenance of volumetric fill influences tissue quality^8^. To build on this relationship, we incorporated repair tissue biomechanics via bubble plot, with larger circles corresponding to a greater equilibrium modulus. The largest circles localized to the upper right quadrant, depicting a zone of “ideal cartilage repair”. Finally, since we observed varying glycosaminoglycan (GAG) loss in the surrounding cartilage (**Figure 1B**), we integrated the Safranin O red intensity of this cartilage as bubble color, revealing that defects in this ideal repair zone provided the best protection. However, we emphasize that most defects remained outside of this zone, highlighting the need to improve the consistency at which defects move into this ideal repair zone.

**Figure 1.**
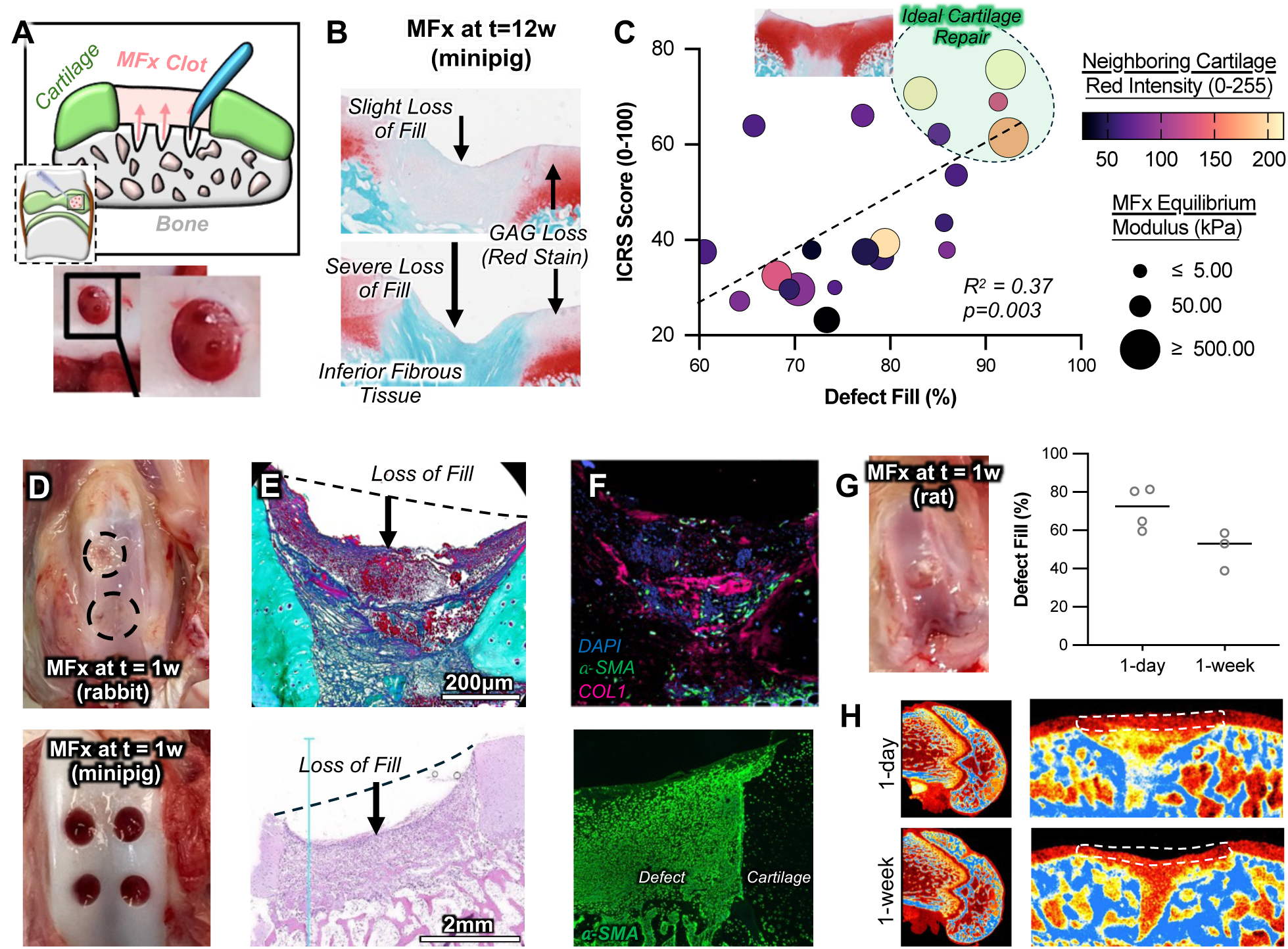
MFx Contraction and Fibrosis. [**A**] MFx procedure involves puncturing subchondral bone to recruit marrow elements. [**B**] Representative MFx defects from minipigs (t=12 weeks), showing two examples of contraction, inferior fibrous tissue formation, and neighboring cartilage GAG loss. [**C**] Histological tissue quality (ICRS Score; 0-100, worst-best) versus defect fill (%), with bubble plot showing repair tissue mechanics with bubble size and neighboring cartilage red staining with color. n=21. [**D**] Rabbit (top) and minipig (bottom) MFx at t=1 week, stained with [**E**] Gomori’s Trichrome (rabbit) or Hematoxylin & Eosin (minipig), and [**F**] DAPI (nuclei), α-SMA, and type-I collagen (COL-1). [**G**] MFx repair in rat model quantified for defect fill with [**H**] representative μCT images at t=1 day and t=1 week. n=3-4.

To investigate early MFx defect fill and repair tissue quality, we observed MFx repair in both a rabbit and pig at one week after surgery (**Figure 1D**). There was clear contraction of the MFx environment (**Figure 1E**), accompanied by fibrotic tissue deposition in the defect area, visualized using immunofluorescence staining for α-smooth muscle actin (α-SMA) and type-I collagen (**Figure 1F**). These findings reinforce the previously defined relationship between defect fill and repair tissue quality and provide evidence that contraction and fibrosis of the MFx defect occur as early as one week. To further explore the temporal reduction in MFx defect fill, we performed MFx in a rat model at 1-day and 7-days (**Figure 1G**). MFx defect fill, quantified using μCT, showed a decrease from 1-day to 7-days post-MFx (**Figure 1H**), verifying that MFx leads to adequate filling initially, but that the provisional clot contracts over time. Histological staining of the rat MFx defects further confirmed the fibrotic susceptibility over the first week (**Figure S2**). Thus, there is motivation to address these early manifestations to drive MFx repair sites more consistently and effectively into this zone of ideal cartilage repair. We believe that MFx augmentations will have a higher likelihood of success if they prevent early volumetric loss and preferentially promote functional cartilage-specific matrix over fibrous scar tissue, enabling protection of the repair site, the neighboring cartilage, and entire joint from long-term wear^7,32^.

### Modulation of initial MFx clot structure may limit contraction but does not improve chondro-fibro ratio

The early MFx repair site is a cellular, fibrin-rich network, formed by the cleavage of fibrinogen by the enzyme thrombin ^33,34^. We visualized this repair environment in rats at one-hour post-MFx surgery using SEM (**Figure 2A**), then simulated this environment *in vitro* using a fibrin gel system (**Figure 2B**), using a combination of fibrinogen, thrombin, CaCl_2_ with embedded MDCs. First, we investigated how altering initial gel properties (fibrinogen/thrombin concentration) impacts contraction and fibrosis. Greater fibrinogen concentration enhances fibrin network formation and overall stability^35,36^; therefore, we hypothesized that added fibrinogen within our fibrin gel would prevent contraction. Nanoindentation confirmed that the effective Young’s Modulus of the gel increased with greater fibrinogen (**Figure 2C**; 10-40 mg/mL), as expected from prior literature ^37^. To investigate early cellular response to the change in mechanics, we fabricated fibrin microgels with encapsulated non-sorted MDCs, to recapitulate the heterogenous milieu of cells recruited to the MFx site^5^. Higher concentrations of fibrinogen led to a rounder cell morphology (similar to cartilage chondrocytes) compared to the irregular, spread-out morphology of cells within low-fibrinogen gels (**Figure 2D**), verified by alterations in cell area and form factor (circularity) (**Figure 2E/F**). Gel contraction was mapped over the first week, with high fibrinogen gels maintaining their area the best (**Figure 2G/H**), comparable to other biomaterial studies that demonstrated less contraction at higher concentrations/mechanics, and suggesting that fibrinogen augmentation of the MFx clot may promote volumetric repair. However, high-fibrinogen gels expressed greater Acta-2 and less Aggrecan (ACAN) expression (**Figure 2I**) leading to a lower ratio of ACAN:Acta-2 (**Figure 2J**), meaning fibrinogen addition may exacerbate fibrotic remodeling.

**Figure 2.**
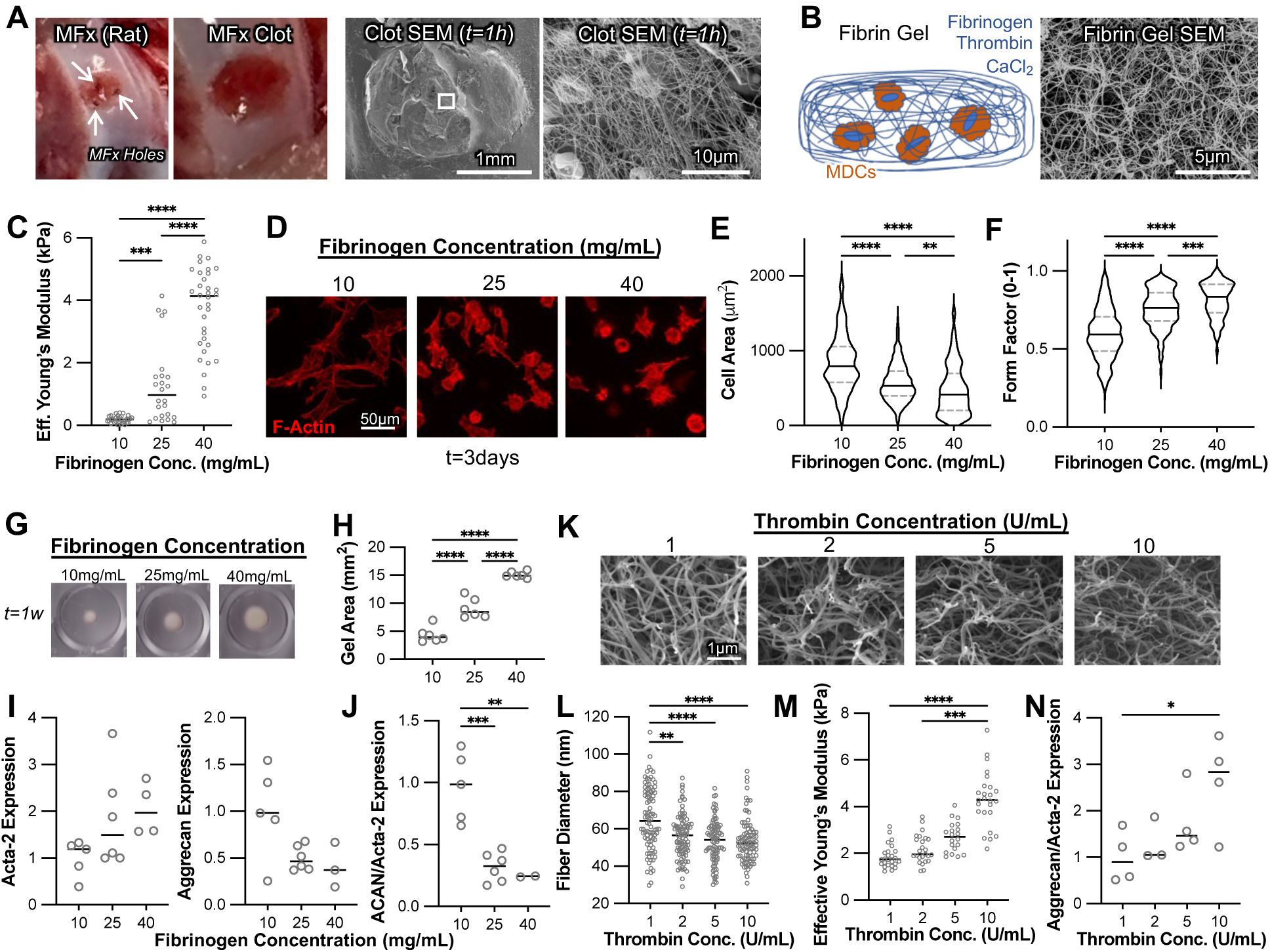
Clot Structure Augmentation. [**A**] Macro-scale images of rat MFx, resulting clot, and scanning electron micrograph (SEM) image at 1-hour post-op. [**B**] Fibrin gel system with associated SEM image that recapitulates MFx clot. [**C**] Effective Young’s Modulus of fibrin gels of varying fibrinogen concentrations, measured via nanoindentation. n>24 indentations/group. [**D**] Representative MDC images in fibrin gels of varying fibrinogen concentrations, and quantified [**E**] cell area and [**F**] form factor (circularity). t=3 days. n>60 cells per group. [**G**] Representative images and [**H**] gel areas at t=1w of culture with varying fibrinogen (10, 25, 40 mg/mL). n=5-6 per group. [**I**] Acta-2 and Aggrecan expression. n=4-6 per group. [**J**] ACAN/Acta-2 Expression. [**K**] SEM Images of thrombin-dependent gels (1-10 U/mL), with [**L**] quantified fiber diameter and [M] Effective Young’s modulus (n>20 indentations/group). [**N**] Aggrecan/Acta-2 expression. n=4 per group. *p<0.05, **p<0.01, ***p<0.001, ****p<0.0001.

Next, we modulated thrombin concentration, with prior evidence of its influence on network architecture^36,38,39^. Similar to these prior studies, higher thrombin concentration led to a denser network of thinner fibrin fibers (**Figure 2K**), verified by quantification of fiber diameter (**Figure 2L**) and led to an appreciable increase in fibrin gel mechanical properties (**Figure 2M**). Interestingly, gels cultured for one week did not exhibit considerable differences in area (similar contraction; **Figure S3A**), but the higher thrombin concentration (10 U/mL) led to the greatest Aggrecan:Acta-2 ratio (**Figure 2N**). This was mostly attributed to decreases in Acta-2 (**Figure S3B**), suggesting a decreased fibrotic phenotype. This was further verified by decreased PAI-1 gene expression^40^ (**Figure S3C**). Thus, modulation of the initial structure via thrombin, but not fibrinogen, demonstrated the greatest promise for volumetric and precise (chondrogenic:fibrotic) repair. Subsequent fibrin gel assays were performed with 25mg/mL fibrinogen and 5U/mL of thrombin to represent an intermediate formulation to study perturbations.

### Extrinsic control of fibrin remodeling prevents contraction but exacerbates fibrosis

Beyond initial structure, the field has demonstrated that governing the rate of provisional matrix remodeling is influential in cell differentiation^41,42^. One central signaling molecule in this process is transforming growth factor beta 3 (TGF-β3), which is critical to endogenous healing. Furthermore, it is the most commonly used growth factor in chondrogenesis studies, previously improving the expression of chondrocyte markers and cartilage matrix deposition^43,44^. While we verified that our MDC’s indeed possessed chondrogenic potential in pellet cultures (Safranin O staining, COL-2 expression, **Figure S4A/B**), TGF-β3 and its other analogs^43,45^ have also been implicated in joint fibrosis^46–48^. To understand the impact of TGF-β3 on MDCs, we stained cells in a monolayer for α-SMA, a myofibroblast marker encoded by the Acta-2 gene. TGF-β3 induced clear α-SMA stress fiber formation and co-localization with F-Actin without changes in cell area (**Figure S4C-E**), indicating myofibroblast propensity with TGF-β3 treatment (verified with Acta-2 gene expression; **Figure S4E/F**), which has been observed in several cell types^49,50^. Next, we explored the addition of TGF-β3 to cells in the context of a 3D fibrin matrix. Interestingly, fibrin gels seeded with MDCs and cultured with TGF-β3 rapidly contracted with clear fibrin degradation (**Figure 3A**). Notably, gene expression of PLAU, which converts plasminogen to plasmin to accelerate fibrinolysis^39^, and its receptor (PLAUR) were significantly increased in TGF-β3-treated gels (**Figure 3B**), as was Acta-2 expression (**Figure 3C**). Thus, while TGF-β3 is certainly chondrogenic, it exacerbates MDC-mediated contraction in a deformable, fibrin-rich environment and simultaneously activates early fibrosis.

**Figure 3.**
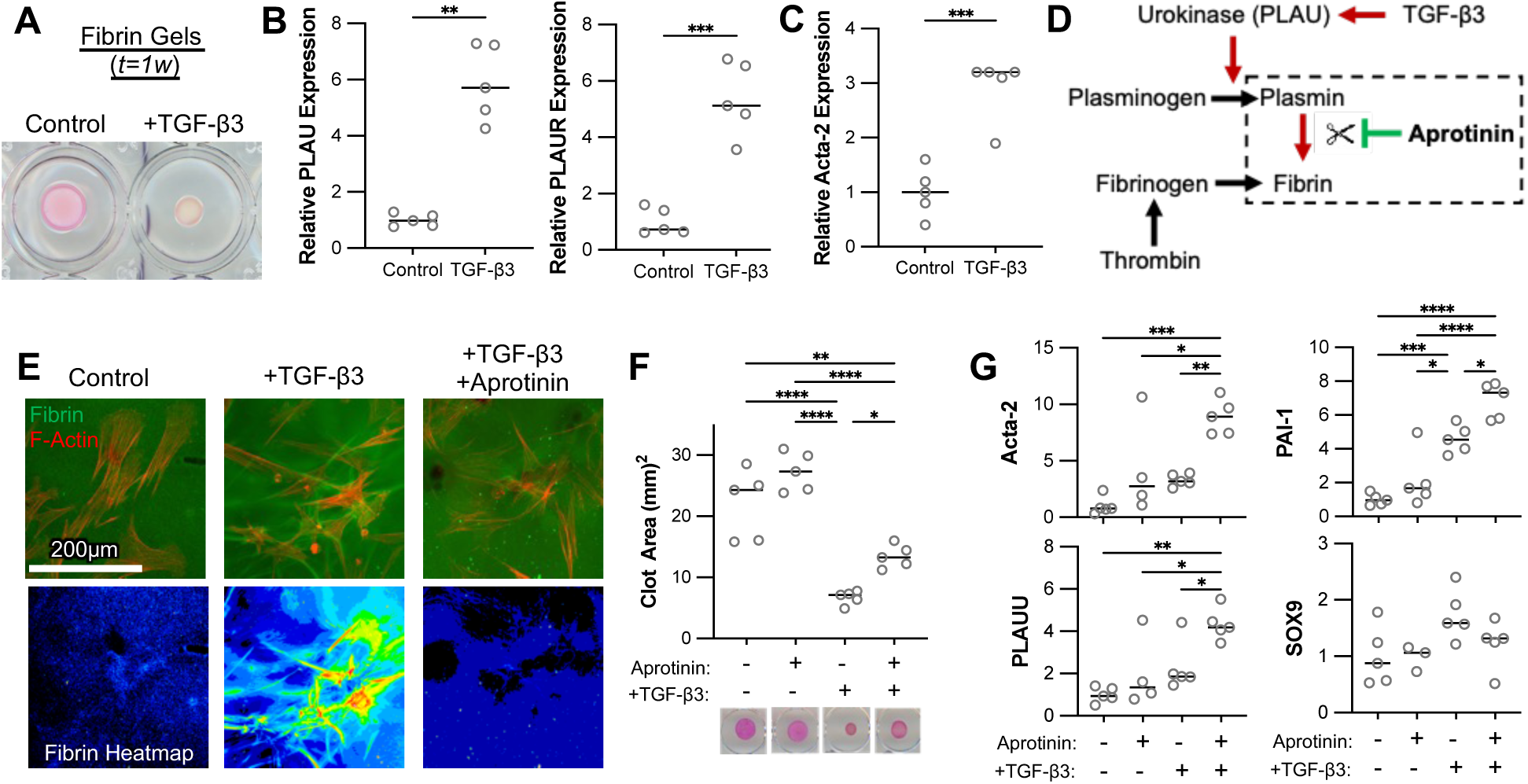
Interplay Between TGF-β3 and Fibrinolysis Prevention. [**A**] Fibrin macrogels treated with control or TGF-β3 media, with [**B**] relative PLAU and PLAUR expression and [**C**] Acta-2 expression. n=5. [**D**] Fibrinolysis pathway and aprotinin activity. [**E**] Confocal images of MDCs and fluorescent fibrin remodeling (t=3 days), cultured +/- TGF-β3 +/- aprotinin. [**F**] Gel area and [**G**] Gene expression (Acta-2, PAI-1, PLAU, SOX9) of TGF-β3 +/- aprotinin treated gels (t=1 week). n=5. *, *p<0.05, **p<0.01, **** p<0.0001.

In an effort to battle the detrimental impacts of TGF-β3, we next sought to reduce extracellular fibrin degradation through the use of aprotinin. Aprotinin is an inhibitor of plasmin (**Figure 3D**), which is a downstream product of plasminogen that initiates the breakdown of fibrin^51^; therefore, we hypothesized that using aprotinin in culture would prevent degradation of the fibrin matrix and potentially prevent fibrosis. We first demonstrated that aprotinin mitigated early TGF-β3-induced fibrin remodeling at the cell-scale, with reduced fibrin densification at t=3 days (**Figure 3E**), observed with fluorescent fibrinogen incorporation into the initial gel. Since fibrin remodeling has been correlated with, and is dependent on, early fibronectin deposition^52^, we performed fibronectin staining of MDCs in fibrin gels cultured with aprotinin and/or TGF-β3 (**Figure S5A**). As expected, TGF-β3 increased fibronectin deposition, but the combination of aprotinin and TGF-β3 reduced deposition to control levels (**Figure S5B**). We next expected early mechano-sensation and cell behavior to track with fibronectin deposition, since the field has routinely linked fibronectin adhesion with mechanosensing. Interestingly, both aprotinin and TGF-β3 increased YAP nuclear localization and exhibited an additive increase in YAP nuclear localization (**Figure S5C**). Given that elevated YAP activity is often a precursor to myofibroblast differentiation and fibrotic matrix production, macro-scale gel studies were performed.

The addition of aprotinin to the culture medium reduced TGF-β3 driven contraction of fibrin clot on a macro-scale and in a dose-dependent manner (**Figure 3F, Figure S6**). However, despite this, aprotinin and TGF-β3 synergistically increased the expression of fibrotic markers Acta-2 and PAI-1 (**Figure 3G**). An increase was also observed in PLAU expression, indicating the cells elevate expression of fibrin-degrading enzymes, but their activity is curtailed, leaving the cells “stuck” in the fibrin gel. Furthermore, no differences in SOX9 expression were observed, meaning that extrinsic control of contraction via fibrinolysis inhibition ultimately decreased the ratio of chondrogenic to fibrotic markers (in this instance exemplified by SOX9 expression: PLAU, PAI-1, and Acta-2 expression), or the “chondro-fibro” ratio. Similar to prior fibrinogen augmentation studies, we acknowledge that fibrin retention at wound healing sites leads to fibrosis^53^, suggesting some degree of fibrin degradation may be necessary to avoid fibrosis. This motivated alternative methods to simultaneously prevent contraction without inhibiting fibrin degradation and enhance cartilage formation.

### Rho-ROCK inhibition synergizes with TGF-β3 to increase early chondro-fibro ratio

To intracellularly alter the ability of cells to contract their surrounding matrix, we chose to modulate cellular contractility via the Rho-ROCK pathway (**Figure 4A**). The Rho-ROCK pathway is involved in various cellular processes, including migration, proliferation, and actin polymerization ^54^. ROCK induces actin reorganization, leading to stress fiber formation and intracellular stress generation^55^. Activation of Rho-ROCK is implicated in fibrosis in several tissues across the body^56,57^; moreover, ROCK inhibitors have been utilized in several applications to reduce contractility, such as for glaucoma and hypertension^58,59^, and have inhibited TGF-β-driven fibrosis in soft tissues (e.g., lung, liver). Notably, TGF-β3 activates the Rho-ROCK pathway, through both non-canonical ALK5 and canonical SMAD signaling ^56,60^. Because of this relationship, several studies have investigated the role of Rho-ROCK in chondrogenesis. ROCK inhibition of cells in a monolayer with the small molecules Y27632 or Fasudil led to increased GAG deposition and SOX9 expression (chondrogenesis) ^61,62^. However, in 3D micromass MSC cultures, ROCK inhibition (without TGF-β3) decreased SOX9 activity and chondrogenesis^63^, though a separate study in micromasses cultured with TGF-β3 resulted in increased chondrogenic gene expression (SOX9, Aggrecan, COL-2) and COL-2 deposition^64^. Conversely, SOX9 activity of chondrocytes in agarose hydrogels was found to be dependent on Rho activity^65^, and ROCK inhibition reduced chondrogenesis. Thus, this interplay between TGF-β3 and the Rho-ROCK pathway exhibits conflicting and context-dependent results, motivating their specific elucidation in MDCs and deformable fibrin-rich systems.

**Figure 4.**
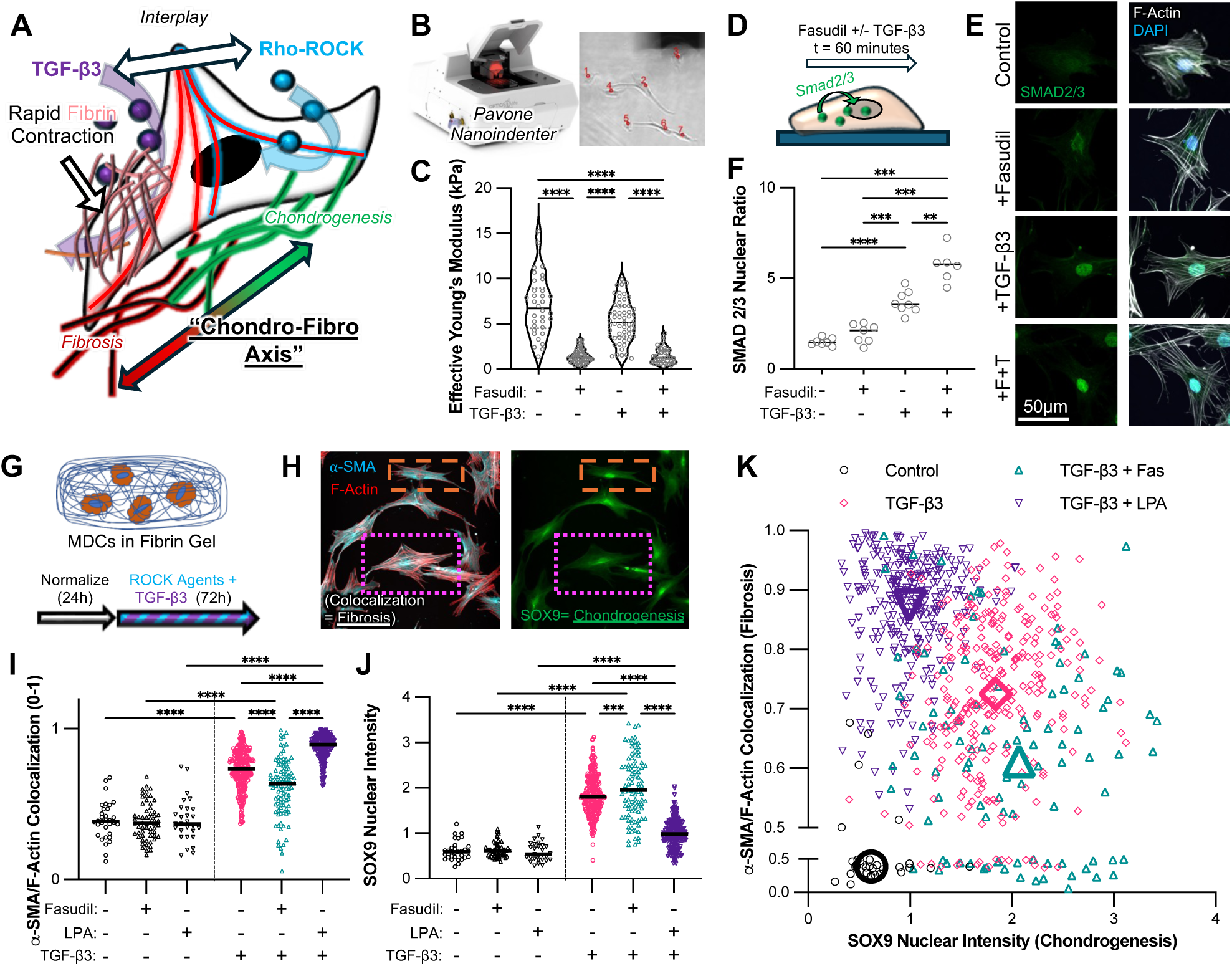
Rho-ROCK Modulation in Conjunction with TGF-β3. [**A**] Schematic of TGF-β3 mediated rapid fibrin contraction, balanced with interplay with ROCK inhibition for activity along chondro-fibro axis. [**B**] Image-guided nanoindentation to obtain [**C**] Effective Young’s modulus of cells treated with Fasudil +/- TGF-β3 for t=1 hour. n>30 cells. [**D**] MDCs in monolayer given Fasudil +/- TGF-β3, and stained for [**E**] SMAD2/3 (t=1 hour), F-actin, and DAPI to enable [**F**] nuclear intensity quantification (n>30 cells). [**G**] MDC in fibrin culture to enable 3-day [**H**] co-staining of α-SMA and SOX9 to quantify chondro-fibro ratio. MDCs treated with combinations of Fasudil, LPA, and TGF-β3 and quantified for [**I**] α- SMA/F-actin co-localization and [**J**] normalized SOX9 nuclear intensity. n>30 cells per group. [**K**] Chondro-fibro axis with α-SMA/F-actin colocalization plotted against normalized SOX9 nuclear intensity. Large symbols represent median value for each group. **p<0.01, ***p<0.001, ****p<0.0001.

To aid in translatability, we utilize Fasudil in our studies, a ROCK inhibitor approved for cerebral vasospasm and pulmonary hypertension in Asia. First, we used image-guided nanoindentation (**Figure 4B**) to verify Fasudil activity, demonstrating reduced MDC stiffness (Effective Young’s Modulus) in only 60 minutes, both in control and TGF-β3 treated cells (**Figure 4C**). Given the ROCK-TGF-β3 interplay, we next elucidated how Fasudil impacts TGF-β3 downstream activation, by quantifying SMAD2/3 nuclear localization at 60 minutes post-treatment (**Figure 4D**). In monolayer, Fasudil led to a modest increase in SMAD2/3 nuclear:cytoplasmic ratio compared to control; this increase was much greater in TGF-β3 treated counterparts (**Figure 4E/F**). Similar enhancement of tenogenic differentiation of MSCs has been observed with Y27632 and TGF-β3 co-treatment^66^. Given the pivotal role of SMAD2/3 in chondrogenesis, these results were promising for augmentation of cartilage repair via combination treatment. Since SMAD2/3 is also a key player in TGF-β induced fibrosis, we also stained monolayer MDCs for α-SMA, demonstrating that Fasudil decreases α-SMA:F-Actin colocalization (**Figure S7**).

We next investigated this interplay between Fasudil and TGF-β3 on early MDC behavior in our deformable, 3D fibrin environment. First, we verified early MDC activity in this system, again showing that Fasudil increases TGF-β3-driven SMAD2/3 nuclear localization (**Figure S8**). However, we then sought to truly acquire an imaging-based metric of both chondrogenesis (SOX9^67–69)^ and fibrosis (α-SMA) to elucidate our chondro-fibro axis on an individual cell basis (**Figure 4G**). Fibrin-MDC gels treated with Fasudil, TGF-β3, and/or lysophosphatidic acid (LPA), an established Rho activator^70,71^. After three days, gels were simultaneously stained for SOX9, DAPI, α- SMA, and F-actin (**Figure S9**). α-SMA stress fiber formation (myofibroblast marker) was measured by calculating the colocalization of α-SMA and F-actin for each cell to obtain an R² value (0–1), and SOX9 expression was quantified by normalizing nuclear SOX9 intensity to DAPI (**Figure 4H**). Neither α-SMA/F-actin colocalization nor SOX9 were affected by the addition of Fasudil or LPA alone (**Figure 4I**, **Figure 4J**). However, TGF-β3 alone increased both α-SMA/F-Actin colocalization and SOX9 nuclear intensity, indicating simultaneous activation of chondrogenic and fibrotic pathways. Co-treatment with TGF-β3 and LPA reduced SOX9 and increased α-SMA, while co-treatment with TGF-β3 and Fasudil resulted in the opposite trend (**Figure 4I**, **Figure 4J**). To visualize both metrics of this chondro-fibro axis simultaneously, we created a correlation plot of F-Actin/α-SMA colocalization versus SOX9 (**Figure 4K**). TGF-β3 clearly moves the median (larger symbol) both upwards and to the right, indicating nonspecific activation of chondrogenic and fibrotic pathways. However, LPA co-treatment leads to a shift towards the fibrotic axis, whereas Fasudil leads to a shift towards the chondrogenic axis, suggesting that Rho-ROCK activity may dictate TGF-β3 activity along a spectrum of early chondrogenesis and fibrosis. These early cell-matrix interactions and subsequent differentiation play a pivotal role in long-term tissue regeneration, giving promise for Fasudil and TGF-β3 co-treatment for more precise chondrogenesis.

### ROCK inhibition eliminates TGF-β3-driven contraction while enabling robust matrix deposition

After establishing these early chondro-fibro axis shifts, we next sought to elucidate the impacts of Fasudil-TGF- β3 treatment on contraction and matrix deposition. First, we performed labeled fibrin gel cultures; after three days, MDCs treated with TGF-β3 alone exhibited strong fibrin densification (**Figure 5A**). However, co-treatment with Fasudil almost completely prevented this densification, with a less contractile phenotype in cells. We next fabricated fibrin macrogels (100 μL, 6 mm diameter), culturing them in media with Fasudil or LPA (10 μM or 50 μM, no TGF-β3) for four weeks. While LPA did not accelerate contraction, it was clear that Fasudil mitigated gel contraction in a dose-dependent manner (**Figure S10A**). Next, we repeated cultures, this time with Fasudil without or with TGF-β3, demonstrating that Fasudil partially mitigated TGF-β3 driven contraction (**Figure S10B**). Excitingly, Fasudil appeared to increase TGF-β3-mediated COL-2 expression (∼200-fold increase over control, **Figure S10C**), further corroborating how early chondrogenic markers (SOX9) previously established with this combination resulted in long-term cartilage-specific matrix expression. We then utilized nascent matrix labeling (via azide-containing analogs) to visualize both nascent protein and nascent GAG deposition in the first week (**Figure S11A**). TGF-β3 clearly increased the integrated intensity of both protein and GAG, while Fasudil co-treatment maintained this TGF-β3 driven level of matrix formation (**Figure S11A/B**).

**Figure 5.**
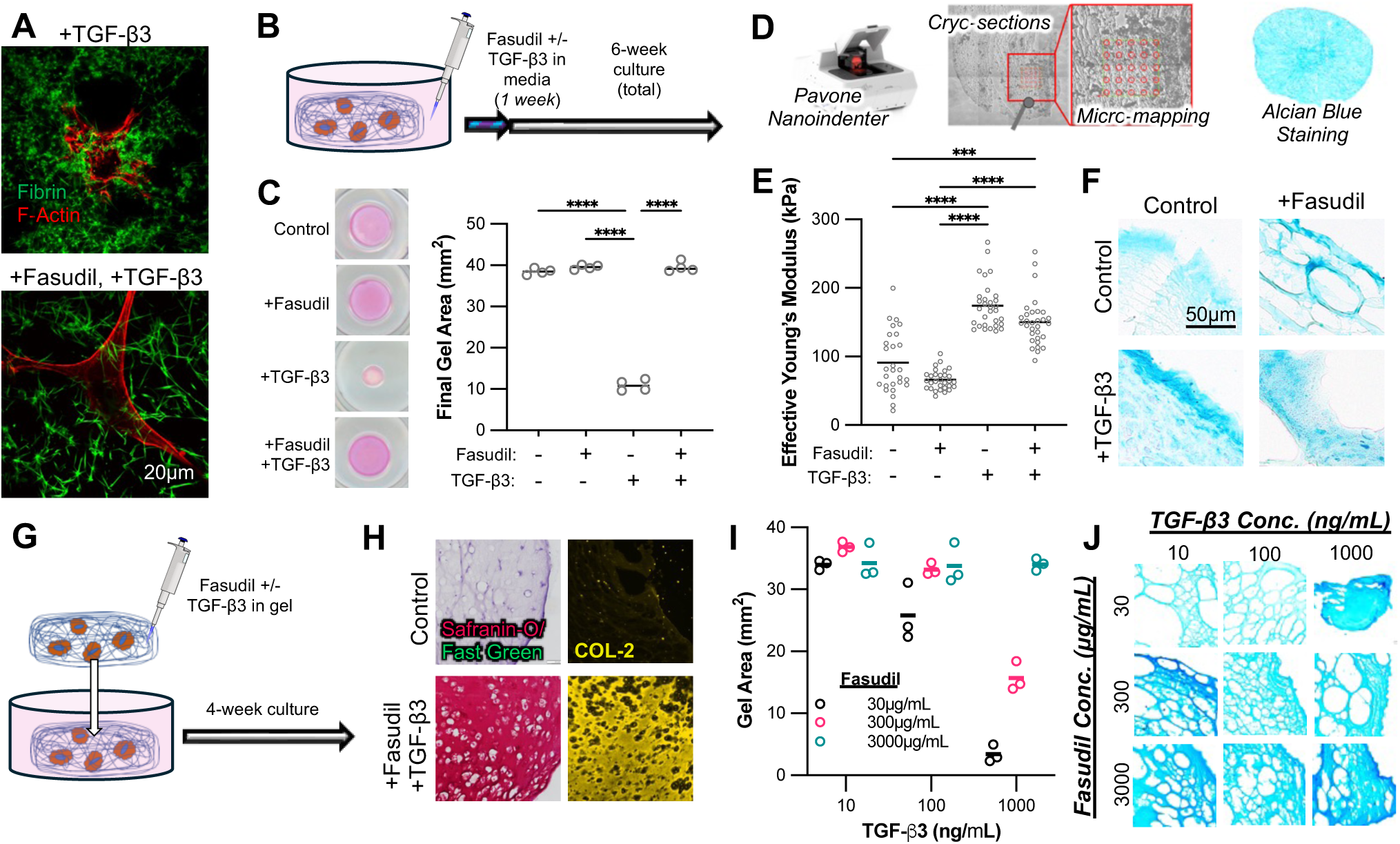
Early Rho-ROCK inhibition impact on long-term behavior. [**A**] MDCs in labeled fibrin gels treated with TGF-β3 or Fasudil+TGF-β3. [**B**] Schematic of 6-week culture with factor application in media for first week. [**C**] Final area of fibrin gels. n=4. [**D**] Image-guided nanoindentation used to obtain [**E**] Effective Young’s Modulus of gel cryosections. n>20 indentations per group. [**F**] Sections stained with Alcian Blue (GAG deposition). [G] Schematic of 4-week culture with factors incorporated into initial fibrin gel. [**H**] Control and Fasudil+TGF-β3 gel sections (t=4w) stained for Safranin-O/Fast Green and type-II collagen (COL-2). [**I**] Final gel area after t=4w of fibrin gels with various doses of Fasuidl and TGF-β3 incorporated into the initial gel. n=3. [**J**] Representative Alcian Blue images from dosing study. ***p<0.001, **** p<0.0001.

Translation into an eventual MFx clot will likely concentrate factor delivery to the early timeframe, rather than exposure for the full four weeks as done previously. Therefore, to mimic expected release over the first week *in vivo*^19^, we conducted a six-week fibrin gel culture, with Fasudil (50 μM) and TGF-β3 (100 ng/mL) added to the culture media only during the first week (**Figure 5B**). Gels treated with TGF-β3 alone contracted significantly compared to control; contraction was entirely mitigated by the inclusion of Fasudil (**Figure 5C**), demonstrating that early drug treatment may be sufficient to prevent longer-term MFx contraction. To verify that Fasudil treatment did not prevent the deposition of functional cartilage-specific matrix, nanoindentation on gel sections (**Figure 5D**) revealed that gels treated with TGF-β3 nearly doubled the Young’s Modulus compared to the controls (**Figure 5E**). This was anticipated, as we hypothesized an increase in matrix density from contraction and anabolic induction of TGF-β3. However, this biomechanical increase was maintained in the gels treated with both TGF-β3 and Fasudil, an increase that is entirely attributed to robust matrix synthesis rather than compaction. These findings were corroborated by Alcian Blue staining to visualize GAG deposition (**Figure 5F**), which demonstrated more robust GAG staining in TGF-β3+Fasudil gels than controls.

To better recapitulate application into the MFx clot, we next incorporated factors into the fibrin gel (30 μg Fasudil, 150 ng TGF-β3) itself prior to formation (**Figure 5G**). After four weeks of culture, combination treatment led to a remarkable increase in GAG and type-II collagen deposition, as visualized by Safranin-O/Fast Green and immunofluorescence staining, respectively (**Figure 5H**). This robust and cartilage-specific maturation was confirmed by gene expression, showing clear increases in aggrecan and type-II collagen expression (**Figure S12A/C**). Although type-1 collagen expression was also increased compared to controls, a significantly elevated type-II to type-I collagen (COL-2/COL-1) ratio was observed in the combination-treated gels (**Figure S12B/D**), which supports an overall promotion of specific chondrogenic potential ^72–74^. Prior to *in vivo* translation, we wanted to investigate the impact of dosing of Fasudil and TGF-β3 within the initial gel on contraction and proteoglycan-rich matrix deposition. We tested three Fasudil concentrations and three TGF-β3 concentrations, identifying a preliminary formulation (300 μg/mL Fasudil, 10 ng/mL TGF-β3) that promotes both volumetric maintenance and robust Alcian Blue staining (**Figure 5I/J**). Together, these results suggest that early, localized co-delivery of TGF-β3 and ROCK inhibition within a fibrin-based gel can effectively direct MDCs toward a stable chondrogenic phenotype without volumetric loss, offering a promising strategy for enhancing cartilage repair in the MFx environment.

### Fasudil + TGF-β3 treatment shows promise *in vivo* as a pharmaceutical augmentation to MFx

Finally, we performed a pilot *in vivo* study to assess the therapeutic potential of Fasudil+TGF-β3 supplementation in MFx-mediated cartilage repair. In Lewis rats, full-thickness chondral defects (2mm diameter) were created in the trochlear groove^75,76^, followed by marrow access (MFx) with a needle to promote marrow bleeding (**Figure 6A**). Once defects were filled with liquid marrow but prior to solidification, a solution of thrombin, CaCl_2_, PBS (control), and Fasudil (1 μg) and/or TGF-β3 (5 ng) was mixed into the clotting marrow using a micropipette (**Figure 6B**). To evaluate the quality of repair tissue and the surrounding cartilage at eight weeks post-MFx, we performed biomechanical mapping within and around the defect (**Figure 6C**) and calculated the equilibrium modulus. While no significant differences were observed between groups in the mechanics of the defect repair tissue (**Figure 6D**), the modulus of the neighboring cartilage was best protected in rats that were treated with both TGF-β3 and Fasudil (**Figure 6E**). Finally, histological analysis revealed that combination treatment led to the greatest deposition of proteoglycan (Alcian Blue staining; **Figure 6F**). These findings suggest that, although combination treatment did not significantly improve the mechanical properties of the defect itself, it may improve tissue quality and lend protection to the region of cartilage surrounding the repair. Further studies will incorporate these factors into a scaffold or drug-delivery microcapsules^19^ to enhance their delivery to the MFx site and ultimately translate this approach towards the clinic.

**Figure 6.**
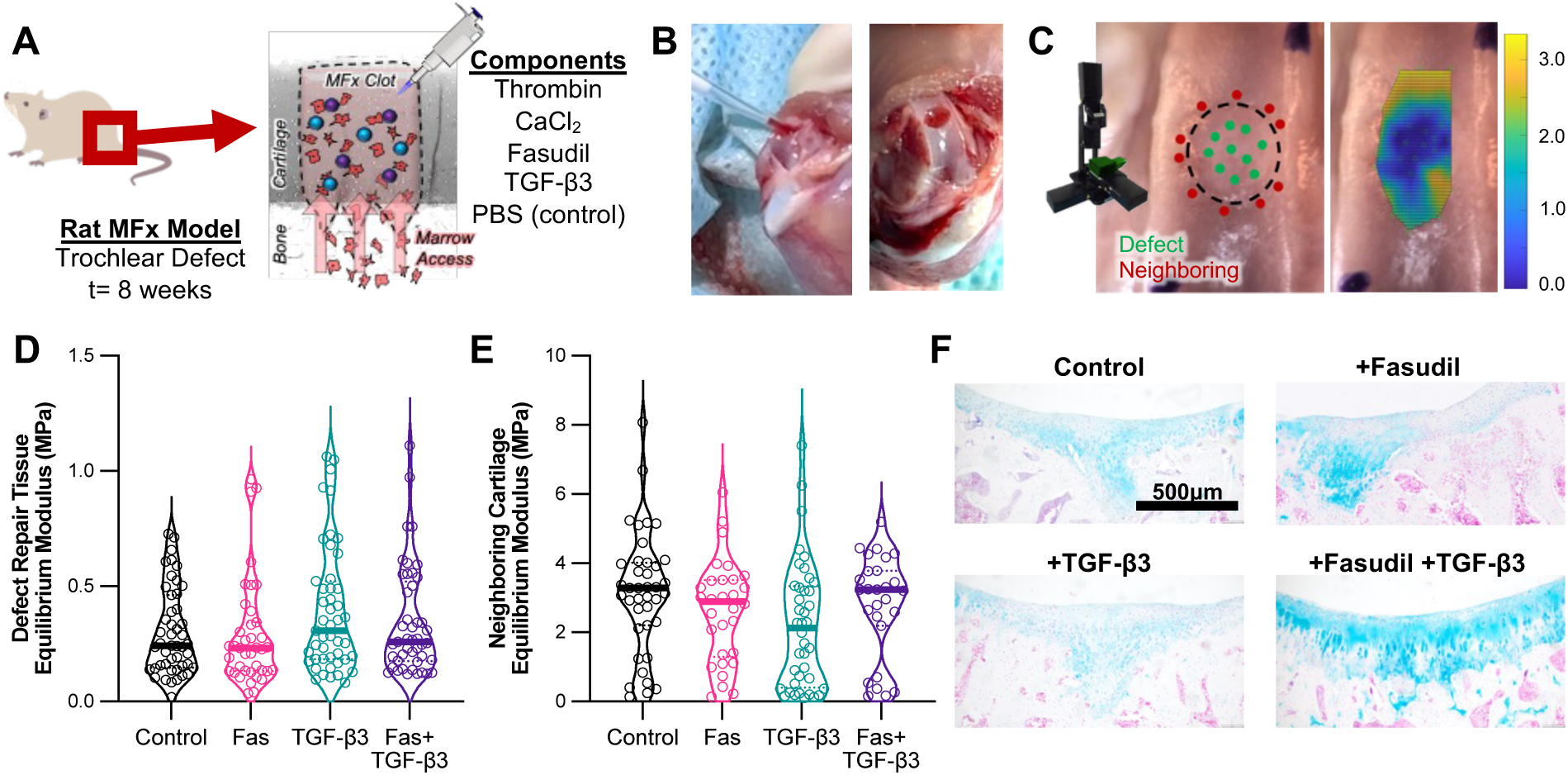
TGF-β3-Fasudil Treatment in Rat MFx Model. [**A**] Schematic representation of treatment in rat trochlear defect model (t=8w, n = 4). [**B**] Incorporation of factors into MFx clot. [**C**] Biomechanical mapping of defect and neighboring cartilage to obtain map across interface. [**D**] Defect and [**E**] neighboring cartilage equilibrium modulus of Control, Fasudil, TGF-β3, and Fasudil+TGF-β3 treated defects. *, **, **** represent p<0.05, 0.01, 0.0001. [**F**] Alcian Blue, Nuclear Fast Red-stained sections.

## Discussion

MFx remains a predominant approach for cartilage repair due to its relative simplicity and affordability. However, the quality and longevity of the tissue regenerated through MFx remain suboptimal, primarily due to inferior fibrous tissue formation and loss of volumetric fill. Our study sought to better understand the biological underpinnings of MFx failure and to develop a targeted strategy to enhance MFx outcomes. We focused on guiding microenvironmental interactions by modulating clot structure, the rate of matrix remodeling, and cell contractility, evaluating both extracellular and intracellular approaches to control contraction. We found that extracellular mitigation of contraction, through fibrinogen augmentation or anti-fibrinolytic treatment, exacerbated fibrosis and pushed cells away from chondrogenesis. Conversely, intracellular control of contraction via the Rho-ROCK pathway, synergizes with TGF-β3 signaling to shift MDCs along the chondro-fibro axis towards a more chondrogenic fate, leading to greater volumetric and cartilage-specific regeneration *in vitro*. These effects were culture-system dependent, as our results in both monolayers and fibrin gels indicate that the context (2D vs 3D, pellet vs fibrin) is critical to this signaling interplay; this study is the first to evaluate this in a deformable fibrin-rich matrix that more accurately represents the early MFx healing environment. Initial pilot data from the rat study including protection of surrounding cartilage and improved proteoglycan deposition gives promise to utilizing these cell-scale mechanobiological findings to improve cartilage repair. Overall, this study lays the foundation for researchers to consider the contractile state and microenvironmental features that direct early cell differentiation and eventual regeneration. We hope to incorporate these factors into a more robust therapeutic strategy, tuning their temporal release and evaluating it as a point-of-care MFx augmentation approach in large animals that better recapitulate human clinical use.

## Methods

### Study Design

The study was designed to investigate the MFx healing environment, modeling the microenvironment *in vitro* to study how cell-material interactions governed matrix remodeling and differentiation. Our first step was to reanalyze data from a prior *in vivo* study in minipigs, followed by short-term studies in one minipig, one rabbit, and rats to elucidate the early microenvironment. All animal studies were conducted under a protocol approved by the Institutional Animal Care and Use Committee. Next, we simulated the MFx environment with a fibrin gel system with encapsulated MDCs, perturbing the environment through structural components, inhibition of fibrin remodeling, growth factor addition, or cell contractility agonists/antagonists, ranging in time point for 60 minutes to 6 weeks (detailed below). *In vivo* pilot therapeutic studies were conducted in skeletally mature male rats (2 months old), with a minimum of four animals per treatment group. Only male rats were included, and the number of animals used is indicated in the figure legends. No animals were excluded from the data analysis*. In vitro* experiments used marrow-derived cells harvested from bovine femoral condyles from three separate donors.

### Animal MFx Characterization Studies

#### In vivo pig model

The 12-week minipig study was previously performed, testing an electrospun hyaluronic acid scaffold containing growth factors^17^. Briefly, we created full-thickness trochlear defects, followed by either MFx or MFx with a TGF- β3 releasing scaffold. Animals were euthanized at 12 weeks, and defects were evaluated using multi-modal outcome measures, including mechanical testing (10-minute stress relaxation with 0.2 mm indentation to yield equilibrium modulus) and histological analysis (Safranin O/Fast Green) for ICRS II histological scoring. We reanalyzed images from that dataset, using histology images to quantify 2D defect fill and red intensity of the surrounding cartilage, combined with ICRS II histological score and equilibrium modulus values from the prior manuscript, to generate the four-variable dataset. One additional Yucatan minipig was included as part of this study in addition to this past cohort for evaluation of early MFx healing. Through a medial patellar arthrotomy, the trochlear groove was accessed. Four 4-mm full thickness cartilage defects were created and subjected to three MFx holes in the subchondral bone with a surgical awl. Defects were allowed to fill with marrow and clot. This one animal was recovered and euthanized at one week post-operatively. Defects were processed for histological and immunofluorescence staining.

#### In vivo rabbit model

To investigate the early MFx environment, we performed MFx in one New Zealand white rabbit. Briefly, through a medial patellar arthrotomy, we laterally dislocated the patella. We introduced two 2mm diameter defects into the trochlear groove, followed by marrow access with a surgical awl. This animal was recovered and euthanized at one week. Osteochondral blocks containing defects were fixed, decalcified, and processed for histology. Paraffin sections were stained with Gomori’s trichrome and immunofluorescence staining was performed for α- SMA and type-I collagen.

#### In vivo rat model

To quantify changes in early defect fill, we performed MFx in 11 male Lewis rats. Lateral dislocation of the patella was used to access the trochlea. Once exposed, a 2mm-diameter flat-ended drill bit was used to create a full-thickness chondral defect in the center of the trochlea. The middle of the defect was then tapped with an 18-gauge needle to enable blood and marrow to flow into the resulting defect and form a clot. The patella was then relocated, and the joint capsule and skin were closed. Postoperative analgesics, antibiotics, and anti-inflammatories were administered according to veterinary guidelines. Rats were euthanized at one hour, one day, and one week (n=4 per group) post-operatively. At the time of euthanasia, joints were harvested and dissected to expose the MFx defect. Samples were subsequently fixed in formalin for 24 hours to prepare for downstream analyses (μCT imaging, scanning electron microscopy, immunofluorescence, and histology).

#### μCT for Defect Fill

The fixed rat MFx samples were debrided of any excess soft tissue and incubated at 37°C in Hexabrix Solution for 30 minutes to facilitate cartilage visualization. Samples were then scanned using a μCT system. The region where the defect was located was contoured and the volume of repair tissue within the entire defect space was calculated to measure the percent defect fill for each sample.

#### Animal Sample Histology

The fixed pig, rabbit, or rat MFx samples were incubated at RT in 14% EDTA decalcification solution for 4 weeks. Decalcification solution was replaced every 3-4 days. Decalcified samples were then placed in 70% ethanol to begin dehydration. Samples were processed through graded ethanol dehydration and xylene clearing before being embedded in hot paraffin wax. Paraffin embedded samples were sectioned at 5 μm at the center of each defect. Samples were stained with Hematoxylin & Eosin, Gomori’s Trichrome, or Safranin-O/Fast Green to visualize early repair tissue. Additional samples were deparaffinized, subject to antigen retrieval (proteinase K), and stained with antibodies for α-SMA and COL-1.

### Tissue Culture Methods

#### MDC Harvest and Culture

Bone marrow-derived cells were harvested from bovine femoral condyles (juvenile, 1-3 weeks old, Research 87). Chunks of bone marrow were excised and rinsed in heparin media (Dulbecco’s Modified Eagle Medium [DMEM] + 2% penicillin-streptomycin-fungizone [PSF] + 0.2% w/v heparin). This solution was then plated and expanded in basal media (BM; DMEM with 10% fetal bovine serum [FBS] and 1% PSF) with media replenishment every 2-3 days. Once cells reached ∼90% confluence, MDCs were trypsinized and frozen in freezing medium (FM; 90% FBS + 10% dimethyl sulfoxide [DMSO]). For experiments, MDCs were thawed and expanded in BM. Bovine MDCs at passages 1-3 were utilized for all *in vitro* experiments.

#### Monolayer Culture

Bovine MDCs were trypsinized, counted, and collected by centrifugation. Cells were then resuspended in BM and seeded into 8-well chamber slides. After 24 hours, the media was replaced with fresh BM containing Fasudil and/or TGF-β3 for study duration (60 minutes to 3 days). Cells were then fixed and processed for immunofluorescence staining.

#### Fibin Micro-gels

To prepare fibrin micro-gels, a fibrinogen solution was mixed with a separate solution containing thrombin, calcium chloride, phosphate-buffered saline (PBS), and the cell suspension. The two solutions were mixed directly within each chamber of an 8-chamber slide to form fibrin gels with final concentrations of fibrinogen (25 mg/mL, unless otherwise noted; Sigma Aldrich F8630), thrombin (5 U/mL, unless otherwise noted; Sigma Aldrich T4648), calcium chloride (20 mM), MDCs (1,750 cells/gel), and PBS to reach a final volume of 10 μL. Chamber slides were incubated at 37°C for 60 minutes to enable gel formation. Basal media (BM; DMEM with 10% fetal bovine serum [FBS] and 1% PSF) supplemented with aprotinin (10 KIU/mL, unless otherwise noted; Sigma Aldrich A1153) was added to the chamber slide, and gels were cultured out to the desired timepoint, with media replenishment every 2-3 days. At the end of the culture, gels were fixed and processed for immunofluorescence. Fluorescently labeled fibrin gels were prepared using the same protocol and concentrations described above, except that a portion of fibrinogen (5% w/w) was replaced with fluorescently labeled fibrinogen (Invitrogen F13191). Gels were protected from light during preparation, incubation, and media changes. Samples were harvested after 3 days of culture. For experiments involving pharmacological treatments, microgels were allowed to normalize for 24 hours in BM before culture media was supplemented with TGF-β3 (10 ng/mL; R&D Systems 243B3), Fasudil (10 μM; Sigma Aldrich CDS021620), and/or LPA (10 μM; Sigma Aldrich L7260) as specified for each experimental group.

#### Fibrin Macro-gels

To prevent fibrin adhesion to well plates, 1% w/v Pluronic solution was added to wells of a 96-well plate and allowed to incubate at RT for 30 minutes. Pluronic was aspirated, and gels were prepared in the treated wells. Unless otherwise specified, fibrin gels were prepared with final concentrations of fibrinogen (25 mg/mL), thrombin (5 U/mL), calcium chloride (20 mM), and MDCs (2 M/mL). Gels were fabricated at a final volume of 100 μL through incubation at 37°C for 60 minutes. Gels were transferred into 24-well plates and cultured in BM + 10 KIU/mL aprotinin, with media replenishment every 2-3 days. At the end of the culture period (7-42 days), gels were harvested for RNA isolation or fixed and sectioned for staining or nanoindentation. To measure contraction, gel area was measured at the start of culture, at every feed, and the terminal timepoint. For experiments involving pharmacologic treatments, gels were allowed to normalize in BM for 24 hours. Results clearly indicate whether pharmacological agents were added throughout culture, for the first week of culture, or within the initial gel.

### Nanoindentation and Electron Microscopy Analysis

#### Micro-gels

Acellular fibrin micro-gels were prepared as previously described. Samples were submerged in PBS to maintain physiological conditions during testing. Nanoindentation was performed using an Optics 11 Pavone nanoindentation system. Indentations consisted of point-indentations across >25 points per gel (27μm radius probe, 0.025N/m). The resulting load-deformation data was analyzed using Optics 11 Pavone analysis software to determine the Effective Young’s Modulus.

#### Monolayer MDCs

MDCs were plated in a monolayer and cultured in BM. After three days, the BM was replaced with fresh BM alone, or BM containing Fasudil (50 μM) and/or TGF-β3 (10 ng/mL) for 45 minutes. Individual cell mechanics were measured using image-guided nanoindentation (3μm radius, 0.025 N/m). The resulting load-deformation data was analyzed using Optics 11 Pavone analysis software to determine the Effective Young’s Modulus.

#### Macro-gel sections

Fibrin macro-gels were snap-frozen in optical cutting temperature (OCT) compound and cryo-sectioned. Samples were rinsed to remove OCT, rehydrated, and submerged in PBS during testing. The gel section was visualized using light microscopy, and grids of indentation points within the gel were tested using an 11 μm radius, 0.5 N/m probe. The resulting load-deformation data was analyzed using Optics 11 Pavone analysis software to determine the Effective Young’s Modulus.

#### Scanning Electron Microscopy

Acellular macro-gels were fabricated with varying fibrinogen and thrombin concentrations, as specified in the Results section. Gels were fixed in 10% Carson’s buffered formalin, rinsed with PBS, and then serially dehydrated from DI H_2_O to 100% ethanol. Samples were then transferred to the Robert P. Apkarian Integrated Electron Microscopy Core at Emory University, where they were underwent snap freezing and cracking to expose the interior of the gel, critical-point drying, and sputter coating. Gels were imaged on the JEOL JSM-IT700HR SEM. Images were binarized in Image J and fiber thickness was calculated using Diameter J.

### Immunofluorescence Staining

Monolayer MDCs or fibrin micro-gels were rinsed in PBS and fixed in 10% Carson’s buffered formalin for 30 minutes. Samples were permeabilized with 1.0% Triton X-100 in PBS for 15 minutes for monolayer samples and 45 minutes for microgels. To reduce non-specific binding, a solution of 3% bovine serum albumin (BSA) was applied for 45 minutes. Primary antibodies, diluted in 1% BSA, were then applied to gels overnight at 4°C. Primaries used included SMAD2/3 (Cell Signaling 5678S; 1:200), α-SMA (Sigma Aldrich A2547, 1:200), SOX9 (ThermoFisher PA5-81966, 1:200), Fibronectin (Sigma Aldrich F6140, 1:200), YAP (ThermoFisher PA5-87568, 1:200). After three PBS rinses, a mouse- or rabbit-secondary antibody (goat Alexa Fluor 488-, 555-, or 647- conjugated, ThermoFisher A11029, A11034, A21424, A21428, A21236, A21245, 1:200 in 1% BSA) was applied for 60 minutes at room temperature. Secondary antibody solution also contained AlexaFluor Phalloidin-546 or - 647 (ThermoFisher A22283, A22287; 1:400). Gels were rinsed and nuclei were stained with Hoechst (ThermoFisher H3570; 1:1000) for 10 minutes. Samples were rinsed and mounted using ProLong Gold mounting medium (ThermoFisher P36934). Cells were imaged using confocal microscopy (Nikon A1R), and images were processed in FIJI for eventual analysis in Cell Profiler. Cell Profiler pipelines were developed to measure cell shape attributes (primarily cell area and form factor), as well as antibody staining features (nuclear intensity, nuclear to cytoplasmic ratio, α-SMA:F-Actin co-localization).

### Nascent Matrix Visualization

#### Media Preparation

Chondrogenic culture medium was prepared, which consisted of DMEM lacking glutamine, L-methionine, and L-cystine, supplemented with 10% FBS, 1% GlutaMAX, 1% PSF, 1% sodium pyruvate, 0.1% ascorbate-2-phosphate, and L-cystine. During media changes, Click-IT™ AHA (L-Azidohomoalanine, Invitrogen, C10102; 0.75 μL/mL) and L-methionine (0.25 μL/mL) were added to the fresh medium. For gels receiving GalNAz treatment, media consisted of DMEM, supplemented with 10% FBS, 1% PSF, 1% sodium pyruvate, and 0.1% ascorbate-2-phosphate. GalNAz (ThermoFisher, 88905; 1 μL/mL) was added to media at each media change. Additionally, gels were treated with TGF-B3 (10 ng/mL) and/or Fasudil (10 μM) throughout the culture period.

#### GelMA Gel Preparation

GelMA gels were prepared by combining GelMA (final concentration of 50 mg/mL), LAP (final concentration of 100 μg/mL), and cell solution (750k cells/mL final concentration) on top of a glass slide. The gel solution was bordered on the top and bottom by two coverslips and a coverslip was added on top of the solution to ensure a flat surface. The gel was exposed to blue light for 5 minutes to crosslink, forming a gel. Next, the coverslips were removed, and the gel was divided into 8 pieces, which were placed into 8-well chamber slides. Gels were rinsed in PBS before adding culture media. Gels were maintained at 37°C and media was replenished every 2-3 days throughout the culture period. After 14 days, the gels were harvested for staining.

### Nascent Matrix Staining

Gels were rinsed in PBS and incubated in 3% BSA for 30 minutes to prevent nonspecific binding. DBCO 488 (20 μL/mL in 1% BSA) was applied to each gel, and gels were protected from light and incubated for 40 minutes. Next, samples were rinsed in PBS and fixed in 10% formalin for 30 minutes. After fixation, CellMask Deep Red (Invitrogen, C10046; 1 μL/mL 1% BSA) was applied for one hour, followed by a 10-minute Hoechst stain (1:1000). Samples were rinsed in PBS and mounted using ProLong Gold mounting medium under a coverslip.

### Analysis of Macro-gels

#### RNA Isolation and qPCR

At the termination of the culture period, fibrin macro-gels were submerged in TRIzol (ThermoFisher #15596018), placed on ice, and homogenized. Chloroform (200μL) was added to each gel solution to initiate phase separation, and the solution was shaken during a five minute incubation at room temperature. Solutions were centrifuged at 12000g for 15 minutes, and the aqueous layer was pipetted into a separate tube. This solution was mixed with equal parts of 70% ethanol, and the resulting solution was added to Bio-Rad Aurum RNA columns, with isolation using the Aurum Total RNA Mini Kit (Bio-Rad #7326820). RNA content and quality was quantified using a Nanodrop. cDNA was synthesized using qScript cDNA SuperMix (QuantaBio #95048-100), generating the same cDNA content for each sample. In 96-well Bio-Rad PCR plates, cDNA was combined with PowerUp SYBR Green qPCR Master Mix (ThermoFisher A25778), DI H_2_O, and primers for β-actin (housekeeping gene) and genes of interest (Acta-2, ACAN, SOX9, COL-1, COL-2, PLAU, PLAUR, PAI-1). qPCR was performed on a Bio-Rad Opus 96 machine and relative gene expression was determined by normalizing output CT values to β-actin and to controls.

#### Histological Preparation

At the termination of the culture period, macro-gels were rinsed in PBS and fixed in 10% Carson’s buffered formalin for two hours. Gels were then moved to plastic molds, submerged in OCT, and allowed to rest at RT for two hours before freezing at -80°C. Gels were sectioned into 10μm sections onto glass slides for nanoindentation (previously described), histology, or immunofluorescence.

#### Histology

Cryosections were rinsed in PBS three times to dissolve residual OCT, followed by two 1-minute rinses in DI H_2_O. For Alcian Blue staining, sections were incubated in 3% acetic acid for 3 minutes before staining with 1% Alcian Blue solution for 30 minutes. For Safranin-O/Fast Green staining, sections were stained with 0.05% Fast Green for 3 minutes, briefly immersed in 1% acetic acid for 10 seconds, and then stained with 0.1% Safranin-O for 15 minutes. Following staining, sections were rinsed with tap water until excess stain was removed. Sections were dehydrated through graded ethanol solutions (70%, 95%, 100%, and 100%) for 2 minutes each, cleared in xylene for 5 minutes, mounted with Permount, and coverslipped. Samples were imaged using light microscopy (Olympus BX63 Upright Microscope).

#### Immunofluorescence

For collagen immunofluorescence staining, cryosections were rinsed with PBS. To induce antigen retrieval, sections were incubated with Proteinase K (20 μg/mL in Tris-EDTA buffer, pH 8.0) for 10 minutes at 37°C. Sections were then allowed to cool to room temperature for 10 minutes and washed twice with PBS containing 0.05% Tween-20. To reduce nonspecific binding, sections were blocked with 3% BSA for 45 minutes. Anti-collagen II (Invitrogen MA5-12789; 1:100) was applied overnight at 4°C in 1% BSA. Sections were rinsed three times with PBS for 5 minutes before incubation with goat anti-mouse Alexa Fluor 555 (1:200 in 1% BSA) for one hour at RT. Sections were washed three times with PBS for five minutes each, counterstained with Hoechst (1:1000) for 10 minutes, rinsed with PBS, and mounted with ProLong Gold mounting medium. Samples were imaged using fluorescence microscopy (Olympus BX63).

### Evaluation of Fasudil/TGF-β3 treatment *in vivo*

#### Rat cartilage repair model

Sixteen male Lewis rats were used for this pilot study under an IACUC-approved protocol. Under isoflurane anesthesia, the left knee of each rat was shaved, cleaned, and surgically draped. Through a medial patellar arthrotomy, lateral dislocation of the patella was used to access the trochlea. Once exposed, a 2mm-diameter flat-ended drill bit was used to create a full-thickness chondral defect in the center of the trochlea. An 18-gauge needle was used as a surgical awl to create one microfracture hole to recruit marrow. Prior to clot formation, a 1 μL of a solution containing thrombin (0.005 U), CaCl_2_ (20 mM), and/or Fasudil (1 μg), TGF-β3 (5 ng), and PBS (amount varied to normalize final volume) was mixed into the marrow-rich blood filling up the defect space using a micropipette. Four groups were tested: Control, Fasudil, TGF-β3, and Fasudil+TGF-β3. Each group received thrombin, CaCl_2_, and PBS. The mixture was allowed to gel before the patella was then relocated, and the joint capsule and skin were closed. Rats were euthanized 8 weeks operatively.

#### Biomechanical Mapping

Rat knee joints were carefully dissected to expose and isolate the MFx defect. Mechanical indentation testing was performed using a Biomomentum Mach-1 multi-axis mapping system. Indentations were conducted within the defect, on the cartilage directly neighboring the defect, and on the healthy cartilage away from the MFx repair site. A 0.5-mm-diameter spherical indenter was used to apply a 20 μm indentation (∼10% of cartilage thickness), which was held for 30 seconds. The resulting data were processed and analyzed in MATLAB using a custom script that determined the instantaneous modulus and equilibrium modulus of the repair tissue and neighboring cartilage^77^. Samples were then fixed in formalin in preparation for histological analysis.

#### Histology

Samples were decalcified in 14% EDTA for 3 weeks, with biweekly solution changes. Following decalcification, samples were dehydrated to 100% ethanol, cleared with xylene, and paraffin-infiltrated. Samples were embedded in paraffin and sectioned at 5μm. Sections were stained for Alcian Blue/Nuclear Fast Red for qualitative analysis of repair tissue quality.

### Statistics

All statistical testing was performed in GraphPad Prism. For comparisons between two groups, unpaired t-tests were used. For analyses of more than two groups, one-way ANOVAs were used, with Tukey’s post-hoc corrections. Outlier and normality testing were performed on individual cell and nanoindentation quantifications (n>10/group). In the event of non-normal data, non-parametric statistical testing was performed. Significance is denoted on graphs with *, **, ***, **** representing p<0.05, 0.01, 0.001, and 0.0001, respectively. Samples sizes are included in figure legends.

## Resource Availability

### Lead contact

Further information and requests for resources and reagents should be directed to and will be fulfilled by the lead contact, Jay Patel.

### Materials availability

This study did not generate new unique materials.

### Data and code availability

The data presented in this work are available from the corresponding author upon reasonable request.

## Acknowledgements

The authors would like to acknowledge the Emory Department of Orthopaedics (Carlos Miller, Hicham Drissi) and the Atlanta VA Medical Center (Colleen Oliver) for their support in completing the proposed work. We also thank Robert Mauck for enabling us to reanalyze data from a prior study. This work was financially supported by internal funds from the Emory Department of Orthopaedics, the Georgia Research Alliance based in Atlanta Georgia under award GRA.25.034.EU.01.b, the Regenerative Engineering and Medicine Center that is supported by the National Center for Advancing Translational Sciences of the National Institutes of Health under Award Number UL1TR002378, a Research Award from the Emory University Research Committee, and the Department of Veterans Affairs (IK2RX003928) and its CReATE Motion Center (I50RX004845).

## Author contributions

Conceptualization, M.H., J.M.P.; Funding Acquisition, J.M.P.; Investigation, M.H., H.S., S.C., A.H., L.M.F., A.Z., W.X.P., N.M.M., A.Y.L.; Methodology, M.H., L.M.F., A.Y.L., N.M.K., J.M.K., J.T.B., J.M.P.; Project Administration, J.M.P., Resources, J.T.B., J.M.P.; Visualization, M.H., H.S., L.M.F., J.M.P.; Writing, M.H., J.M.P.; Review and Editing, All.

