## Supplementary material for "Directing the Chondro-Fibro Axis via Early Microenvironmental Interactions to Enable Precise and Volumetric Cartilage Repair": Figure S

### Supplementary Figures

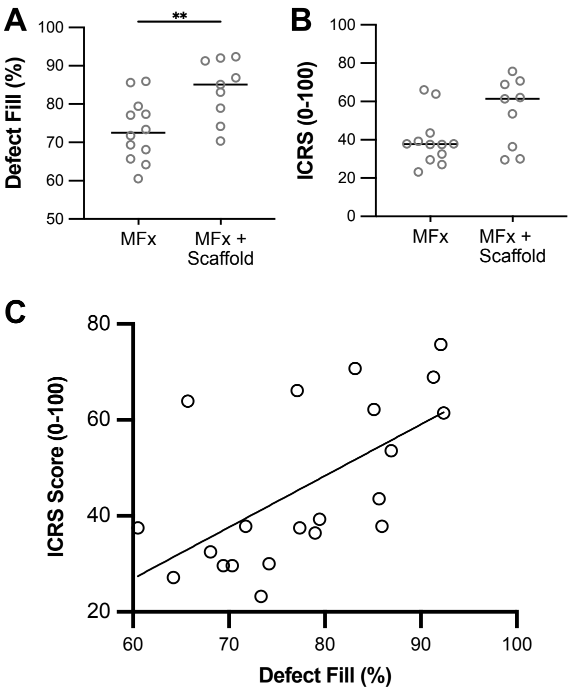


**Figure S1. Yucatan Minipig Defect Fill and ICRS Data.** [**A**] Defect fill (%) and [**B**] ICRS score of defects in MFx and MFx + Scaffold groups. t=12 weeks. n=9-12/group. [**C**] Plot of ICRS score vs Defect fill.

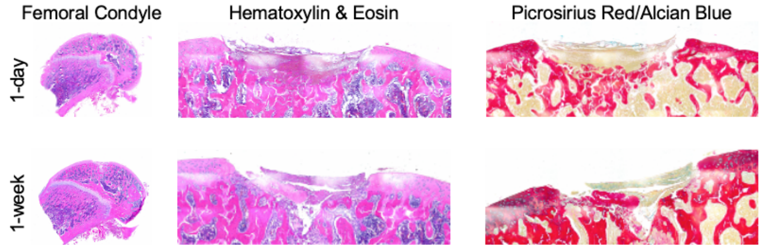


**Figure S2. Rat MFx at Early Time Points**. Histology images (whole femoral condyle and defects) with either Hematoxylin & Eosin or Picrosirius Red/Alcian Blue staining of early rat MFx (t=1-day and 1-week).

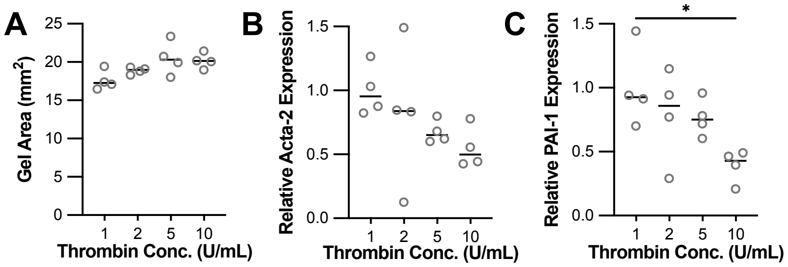


**Figure S3. Thrombin-dependent Macro-gel Contraction and Expression.** [**A**] Gel areas at t=2 weeks of culture with varying thrombin concentration (1, 2, 5, and 10U/mL). [**B**] Gene expression of Acta-2 [**C**] and PAI-1. n=4/group. *p<0.05.

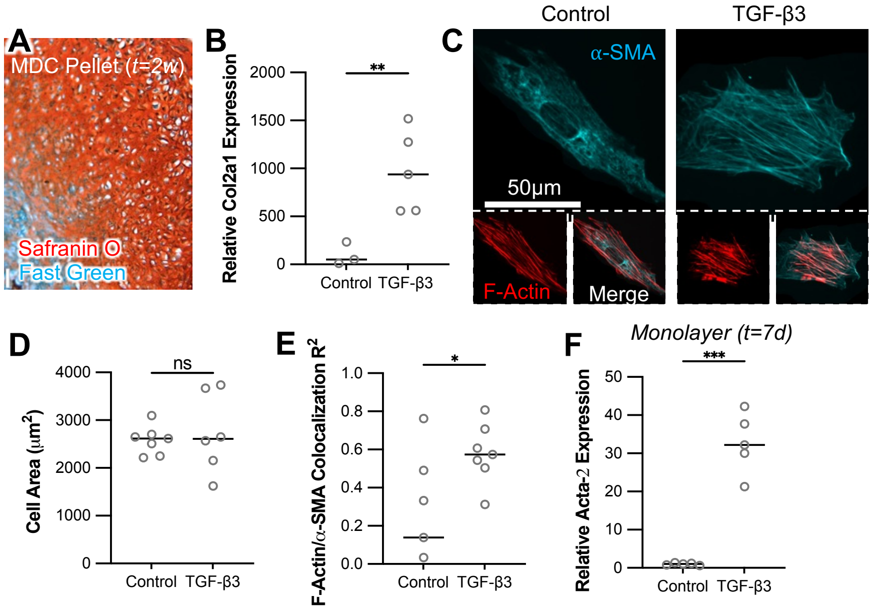


**Figure S4. MDCs undergo TGF-β3-induced chondrogenesis in pellet culture but exhibit fibrosis in monolayer.** [**A**] MDC pellet at t=2 weeks, stained for Safranin-O Fast Green and [**B**] type II collagen expression. n=3-5/group. [**C**] MDCs in monolayer stained for α-SMA and F-actin. t=3 days in culture. Quantification of [**D**] cell area and [**E**] F-actin/α-SMA co-localization. n=5-7 cells/group. [**F**] Gene expression of Acta-2 of MDCs in a monolayer. t=7 days. n=5/group. *p<0.05, **p<0.01, *** p<0.001.

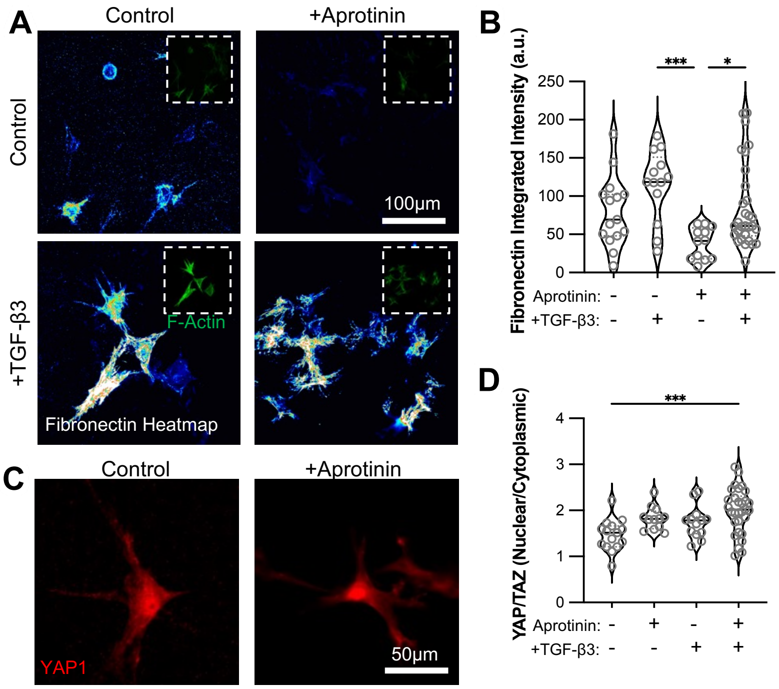


**Figure S5. Combination TGF-β3 and aprotinin treatment reduces fibronectin deposition but enhances mechanosensation.** [**A**] Heatmap of fibronectin staining of MDCs in micro-gels cultured with aprotinin (100KIU/mL) and/or TGF-β3 (10ng/mL). t=3 days. [**B**] Quantification of fibronectin intensity. [**C**] YAP staining of MDCs in micro-gels and [**D**] quantification of YAP nuclear localization. *p<0.05, *** p<0.001.

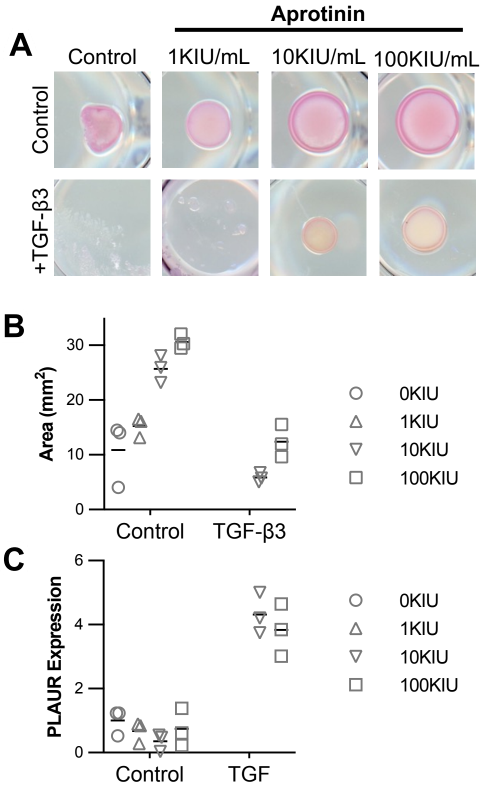


**Figure S6. Aprotinin reduces TGF-β3-driven gel contraction in a dose-dependent manner.** [**A**] Representative images of macro-gels cultured with TGF-β3 (10ng/mL) and/or aprotinin, at the termination of culture period (t=7 days). Quantification of [**B**] gel area and [**C**] PLAUR gene expression. n=3/group.

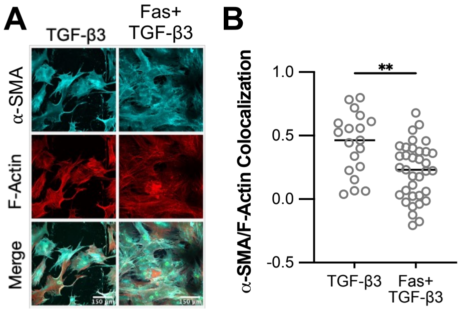


**Figure S7. ROCK inhibition decreases F-actin/α-SMA colocalization in TGF-β3-treated cells.** [**A**] α-SMA and F-actin staining of MDCs in a monolayer treated with TGF-β3 (10ng/mL) and/or Fasudil (50μM) for t=60 minutes. [**B**] Quantification of F-actin/α-SMA co-localization. n>19 cells/group. **p<0.01.

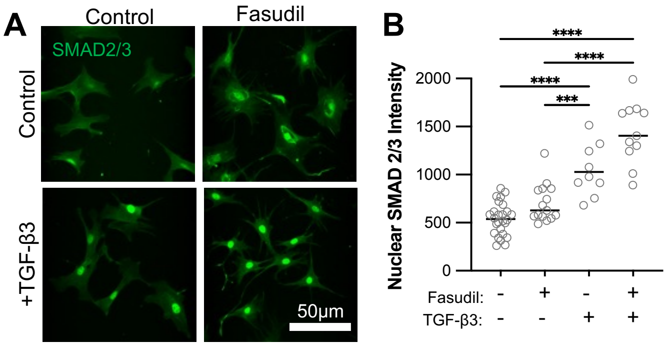


**Figure S8. Rho-ROCK inhibition improves early TGF-β3-driven SMAD2/3 nuclear localization.** [**A**] Cells on glass treated with TGF-β3 (10ng/mL) and/or Fasudil (50μM) for t=60 minutes. [**B**] Quantification of nuclear SMAD2/3 intensity. n>9 cells/group. ***p<0.001, ****p<0.0001.

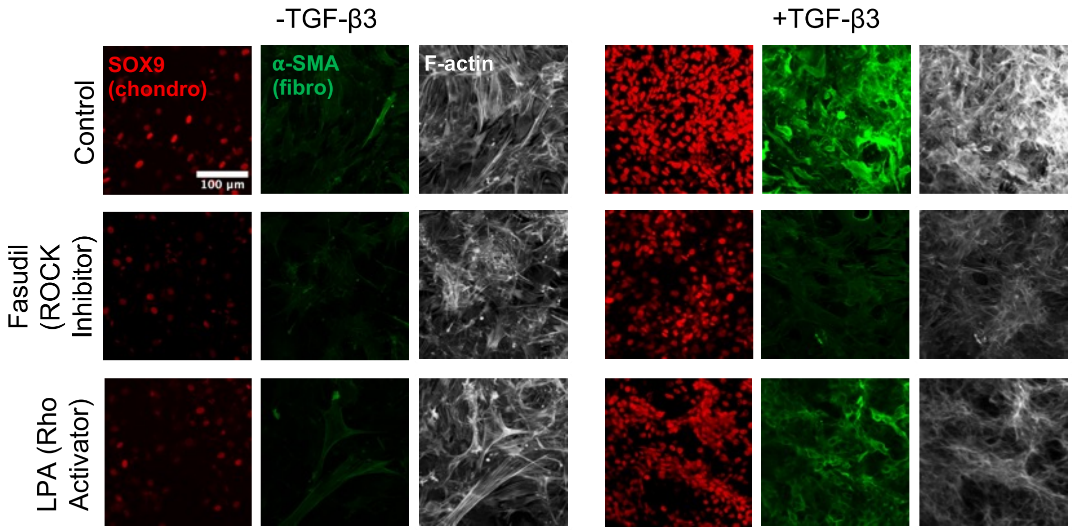


**Figure S9. Rho/ROCK inhibition with TGF treatment enhances SOX9 activation and reduces F-actin/α-SMA co-localization in 3D fibrin gels.** Representative images of MDCs encapsulated in fibrin gels and cultured in BM. ROCK inhibitor (Fasudil) and Rho activator (LPA) were added (10μM) as well as TGF-β3 (10ng/mL) after 2 days. Gels were fixed and stained after t=3 days of treatment.

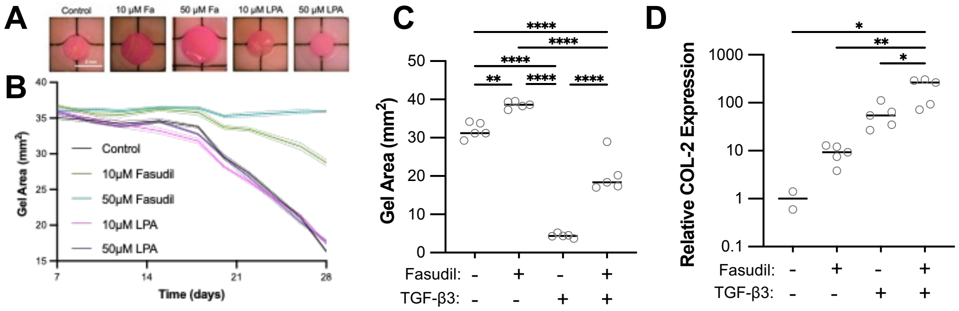


**Figure S10. Fasudil reduces TGF-β3-driven fibrin gel contraction and enhances COL-2 expression.** [**A**] Representative images of macro-gels at the termination of the culture period (t=4 weeks). Gels were cultured in BM supplemented with varying doses of Fasudil and LPA. [**B**] Quantification of macro-gel area throughout the culture period, measured every 2-3 days, demonstrating dose-dependent differences in gel contraction. [**C**] Final gel area of gels cultured in BM supplemented with TGF-β3 (10ng/mL) and/or Fasudil (50µM) after 4 weeks. n=5 gels/group. [**D**] Relative COL-2 gene expression. *p<0.05, **p<0.01, ***p<0.001, ****p<0.0001.

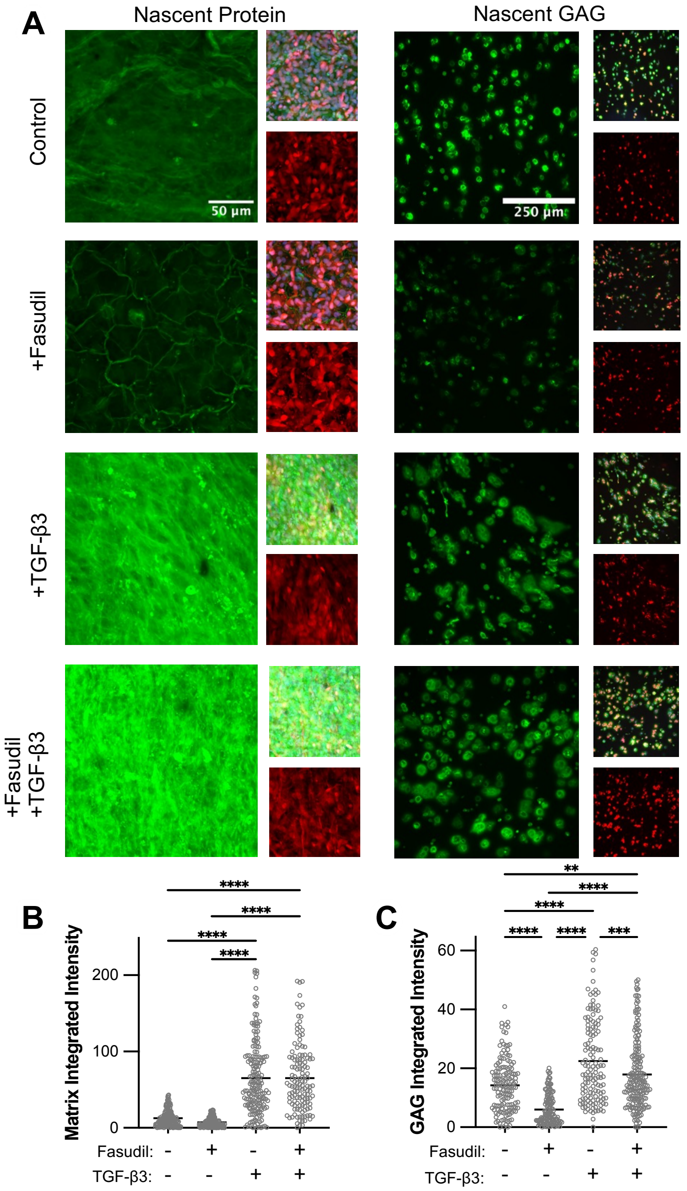


**Figure S11. Fasudil maintains TGF-β3-induced nascent matrix deposition.** [**A**] Representative images of nascent protein and GAG deposition of MDC-laden GelMA gels cultured for t=7 days in BM supplemented with TGF-β3 (10ng/mL) and/or Fasudil (10μM). Newly synthesized proteins and GAGs were metabolically labeled using azide-containing analogs. Green indicates nascent protein or GAG labeling, and red indicates the plasma membrane. [**B**] Quantification of integrated fluorescence intensity for nascent protein labeling. [**C**] Quantification of integrated fluorescence intensity for nascent GAG labeling. n>40 cells/group. **p<0.01, ***p<0.001, ****p<0.0001.

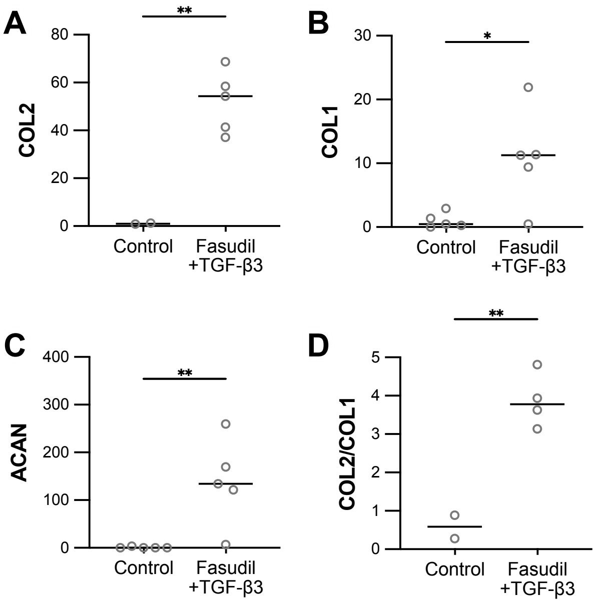


**Figure S12. Gel-incorporated Fasudil and TGF-β3 supports cartilage-specific maturation.** Macro-gels incorporated with Fasudil and TGF-β3 were evaluated for chondrogenic gene expression following t=4 weeks of culture in BM. Relative expression of [**A**] aggrecan (ACAN), [**B**] COL1, [**C**] COL2, and [**D**] COL-2/COL-1 ratio. n=5 per group. $^*p<0.05$, $^{**}p<0.01$.
